# Deficiency of the E3 Ubiquitin Ligase RBCK1 Causes Diffuse Brain Polyglucosan Accumulation and Neurodegeneration

**DOI:** 10.1101/277392

**Authors:** Mitchell A. Sullivan, Felix Nitschke, Erin E. Chown, Laura F. DiGiovanni, Mackenzie Chown, Ami M. Perri, Sharmistha Mitra, Xiaochu Zhao, Cameron A. Ackerley, Lori Israelian, Saija Ahonen, Peixiang Wang, Berge A. Minassian

## Abstract

Glycogen synthesis is vital, malstructure resulting in precipitation and accumulation into neurotoxic polyglucosan bodies (PBs). One well-understood mechanism of PB generation is glycogen branching enzyme deficiency (GBED). Less understood is Lafora disease (LD), resulting from absence of the glycogen phosphatase laforin or the E3 ubiquitin ligase malin, and accumulation of hyperphosphorylated PBs. LD afforded first insight that glycogen sphericity depends on more than adequate branching activity. Unexpectedly, deficiencies of the Linear Ubiquitin Chain Assembly Complex (LUBAC) components RBCK1 and HOIP result in PBs in muscle tissues. Here we analyzed nervous system phenotypes of mice lacking RBCK1 and find profuse PB accumulations in brain and spinal cord with extensive neurodegeneration and neurobehavioral deficits. Brain glycogen in these mice is characterized by long chains and hyperphosphorylation, similar to LD. Like in LD, glycogen synthase and branching enzyme are unaltered. Regional PB distribution mirrors LD and not GBED. Perisynaptic PB localization is unlike LD or GBED. The results indicate that RBCK1 is part of a system supplementing laforin-malin in regulating glycogen architecture including in unique neuronal locales.

## INTRODUCTION

Intracellular storage of blood-derived glucose in the form of highly branched glycogen molecules is critical for life, allowing large amounts of glucosyl residues to be stored in compact yet soluble particles. Crucial to glycogen’s solubility is its relatively short chains and high abundance of evenly distributed branch points, preventing the chains from forming double helices and precipitating (which is the case in amylopectin, the major glucose storage molecule in plants). The principle enzymes involved in glycogen synthesis, glycogenin (GYG1, synthesis initiation), glycogen synthase (GYS1/GYS2, chain elongation) and glycogen branching enzyme (GBE1, branching) have been known for decades^1-3^. GBED (Glycogen Storage Disease IV; GSD IV) leads to a pathological accumulation of poorly branched insoluble glycogen, which precipitates and accumulates into PBs^4^. The age of onset and severity of GSD IV is determined by the amount of residual GBE1 activity. Near total absence of GBE1 results in the infantile Andersen’s disease, a severe condition which leads to childhood liver failure and early fatality^5^. Conversely, mutations associated with higher residual GBE1 activity (most commonly in patients homozygous for the GBE1 p.Y329S variant) result in Adult Polyglucosan Body Disease (APBD)^6^. In these patients, despite the presence of PBs in skeletal muscle, heart and liver, symptoms are limited to the nervous system. In the latter, PBs amass at neuronal axon hillocks and tightly within long axons, which they frequently appear to completely clog. The associated disease consists of upper and lower motor-neuron dysfunction, distal sensory loss, neurogenic bladder and mild cognitive decline^7^.

Since the late 1990’s another system has been shown to be essential to the formation of correctly branched glycogen, namely the complex of laforin (EPM2A), a glycogen phosphatase with a carbohydrate-binding domain^8-11^, and malin (NHLRC1), a RING domain-containing E3 ubiquitin ligase with a growing list of proposed substrates^12-15^. Deficiency of either laforin or malin leads to the accumulation of highly phosphorylated PBs in many glycogen producing tissues, including the brain. Unlike in APBD, PBs in LD accumulate not in neuronal long axons but in their cell bodies and dendrites. This is associated not with an axonopathic disease but with a progressive myoclonic epilepsy with death from increasingly intractable seizures usually within a decade of onset^16,17^.

Most recently, yet another system was implicated, with the absence of the RANBP2-type and C3HC4-type zinc finger containing 1 (RBCK1) protein resulting in the formation of PBs in skeletal and cardiac muscle in both humans and mice^18-20^. RBCK1 (58 kDa, also known as HOIL-1) is best known to form a ∼600 kDa complex with two other proteins, HOIL-1L Interacting Protein (HOIP) and SHANK-associated RH domain-interacting protein (SHARPIN). This complex, LUBAC, mediates M1-linked linear polyubiquitination, a modification that enables nuclear translocation of nuclear factor-кB (NF-кB) and its pleiotropic immune system-critical transcriptional regulation. Both RBCK1 and HOIP contain a RING-between-RING (RBR) domain. However, the LUBAC-mediated ubiquitination is achieved via HOIP’s catalytic domain, with RBCK1’s role in the complex seemingly limited to a critical interaction with HOIP’s auto-inhibitory domain, preventing the inhibition and allowing HOIP to function^21,22^. RBCK1 has also been proposed to target a varied set of proteins for ubiquitination and proteasomal degradation potentially through its own RBR domain, independent of LUBAC^23-27^. However, most of this evidence was acquired through *ex vivo* overexpression experiments and remains unsettled.

Patients and mice deficient of RBCK1 have a unique combination of hyper-inflammation and immune deficiency that appears to variably reach clinical significance from none to fatality based on particular pathogen exposure and other factors. Both patients and mice exhibit progressive PB accumulation in skeletal muscle and heart, culminating, in humans, in skeletal myopathy and heart failure^18,20,28^.

Here we use an RBCK1 knockout (KO) mouse model to explore the role of RBCK1 in glycogen metabolism and PB formation. Unexpectedly, the brain is the predominantly affected organ, with extensive PB accumulations and neurodegeneration. We compare the brain regional distribution and mechanism of formation of these PBs to those in mouse models of LD (*Epm2a^-/-^*) and APBD (*Gbe1*^Y329S/Y329S^) and find that they parallel, though imperfectly, those of LD.

## RESULTS

### Distribution of brain PBs in RBCK1 KO mice closely but incompletely overlaps that of LD

Treatment of tissue sections with periodic acid-Schiff diastase (PASD), allows the staining of glycogen-like material that is relatively resistant to amylase (historic: diastase) degradation, such as PBs^29^. Figure 1 shows the PASD staining of sections from different parts of the brain, spinal cord, skeletal muscle, heart, and liver of wildtype (WT) and RBCK1 KO mice of various ages. There are small levels of PASD stained deposits in the hippocampus of the 14-month-old WT mice, likely corpora amylacea, normally observed in aged humans and mice^30-32^. While it has been reported that these PBs, stained by PAS, are sensitive to diastase digestion in mouse samples^31^, we still observed them here. All other sections of the WT mouse brain were free from visible polyglucosan accumulation.

**Figure 1.**
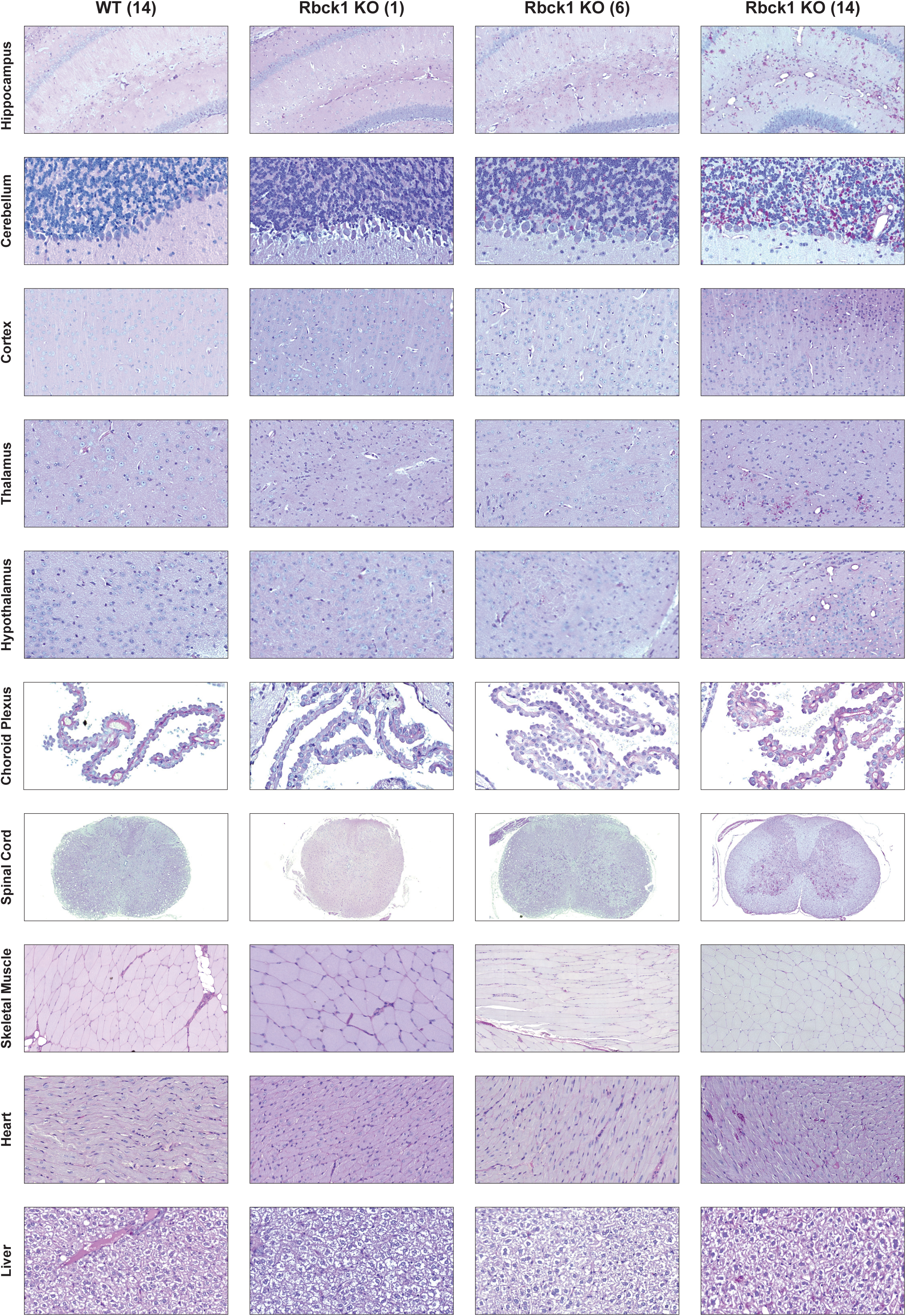
Progressive PB accumulation in RBCK1 KO mouse tissues, including brain regions (PASD stain). Intense pink stains are the PBs that resist digestion with the glycogen degrading enzyme amylase (diastase); age of mice in months indicated in parentheses. A table that summarises the relative amount of accumulation is given in the Supplemental Information, Table S1.

Similar to the mouse models of LD (*Epm2a^-/-^*; referred to henceforth as laforin KO)^33^ and APBD (*Gbe1^Y329S/Y329S^*; referred to henceforth as GBE1-deficient)^34-36^, there are significant deposits of PASD stained material in the brains of RBCK1 KO mice, which increase with age (Figure 1). In the hippocampus, there is a higher concentration of PBs in the dendrite-dense stratum lacunosum-moleculare in RBCK1 KO mice. The polyglucosan accumulation in the laforin KO mice and GBE1-deficient mice have PBs more evenly distributed throughout the hippocampus (Figure 2). The PBs in the RBCK1 KO and laforin KO mice are large in appearance, compared to the smaller ubiquitously spread accumulations in the GBE1-deficient animals.

**Figure 2.**
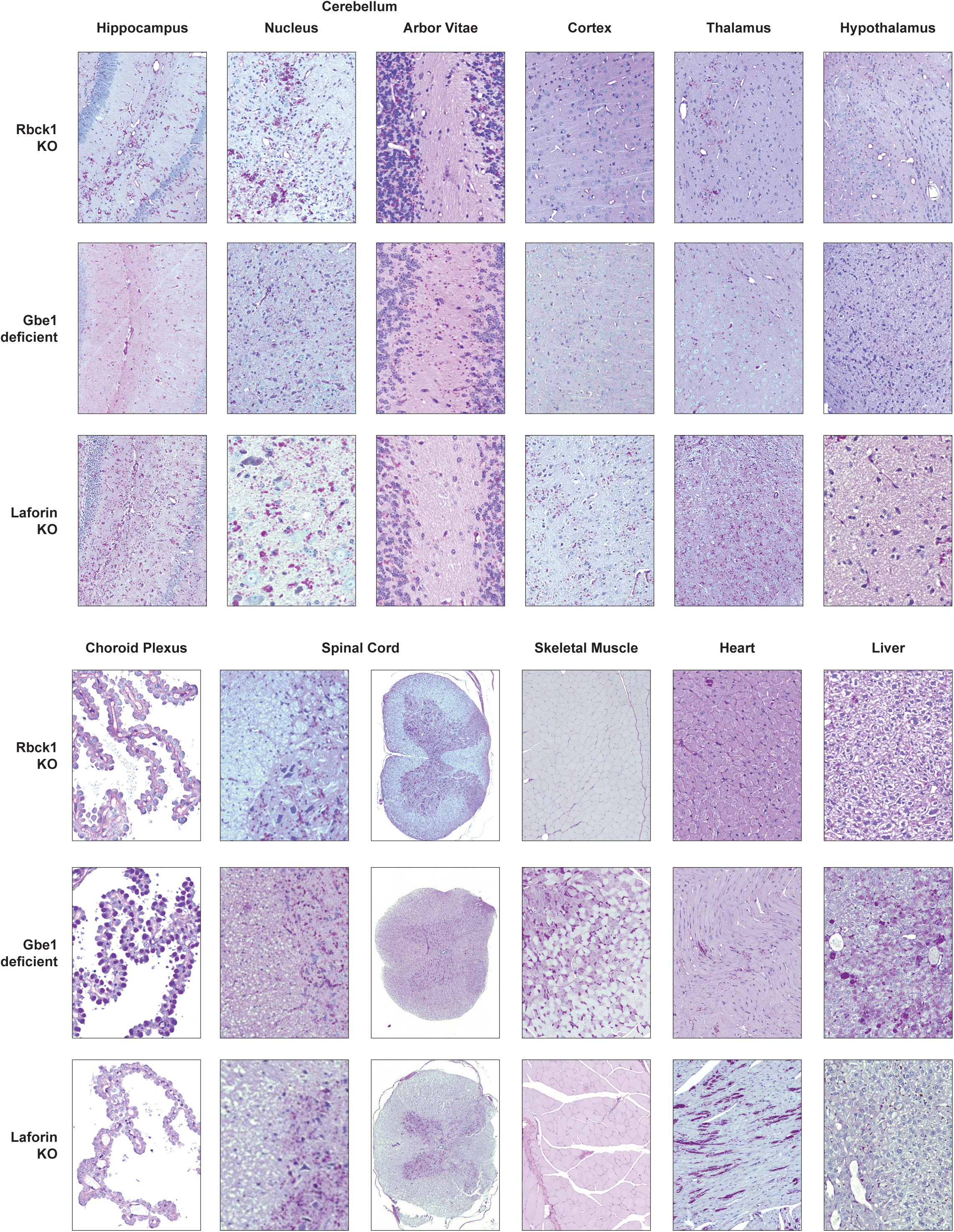
Comparison of PBs in RBCK1 KO, laforin KO and GBE1-deficient mouse tissues and brain regions (PASD stain); all mice 12-14 months of age. A table that summarises the relative amount of accumulation is given in the Supplemental Information, Table S2.

In the cerebellum (Figure 2), laforin KO and RBCK1 KO mice show a high abundance of large PBs throughout the granular layer and the nuclei (dentate, interposed and fastigial), while no evidence of PBs was found in the white matter of the arbor vitae. GBE1-deficient mice, however, have PBs scattered ubiquitously through all areas of the cerebellum, including the arbor vitae.

Another striking difference comparing RBCK1 KO and laforin KO mice with GBE1-deficient mice, is a less consistent accumulation of PBs in the cortex. While in RBCK1 KO and laforin KO there are clusters of PBs, GBE1-deficient mice have small PBs spread ubiquitously throughout the cortex.

Also, while there are clusters of bodies in the thalamus and hypothalamus of RBCK1 KO mice, the PBs were less frequent than in laforin KO and GBE1-deficient mice. Again, the GBE1 deficient mice had a large number of small PBs spread ubiquitously, while the laforin KO and RBCK1 KO mice had PBs concentrated in specific, dendrite-rich areas.

In the choroid plexus, there appear to be no PBs in RBCK1 KO mice, very few if any in laforin KO mice, and massive accumulations in GBE1-deficient mice. Nearly every cell in the choroid plexus of GBE1 deficient mice appears completely occupied by PBs.

In the spinal cord, PBs are found exclusively in the grey matter in both RBCK1 KO and laforin KO mice, while in GBE1 deficient mice the PBs are scattered ubiquitously throughout both the grey and white matter.

As has previously been described in humans and mice^18,37,38^, PBs are seen in the cardiac muscle of RBCK1 KO mice. Compared to human cases, the mouse skeletal muscle contained no significant accumulation of PBs. There were also no bodies in the liver, not inconsistent with human patients, only some of whom have hepatic PB accumulation^18^ (Figure 1 and 2).

The glycogen content of brain, skeletal muscle, heart and liver from WT and RBCK1 KO mice was analyzed (Figure 3A). Glycogen content was increased in RBCK1 KO, only in the brain. Compared to the WT controls 14-month-old RBCK1 KO mice have on average ∼2.6 times higher levels of brain glycogen, consistent with the accumulation of PBs.

**Figure 3.**
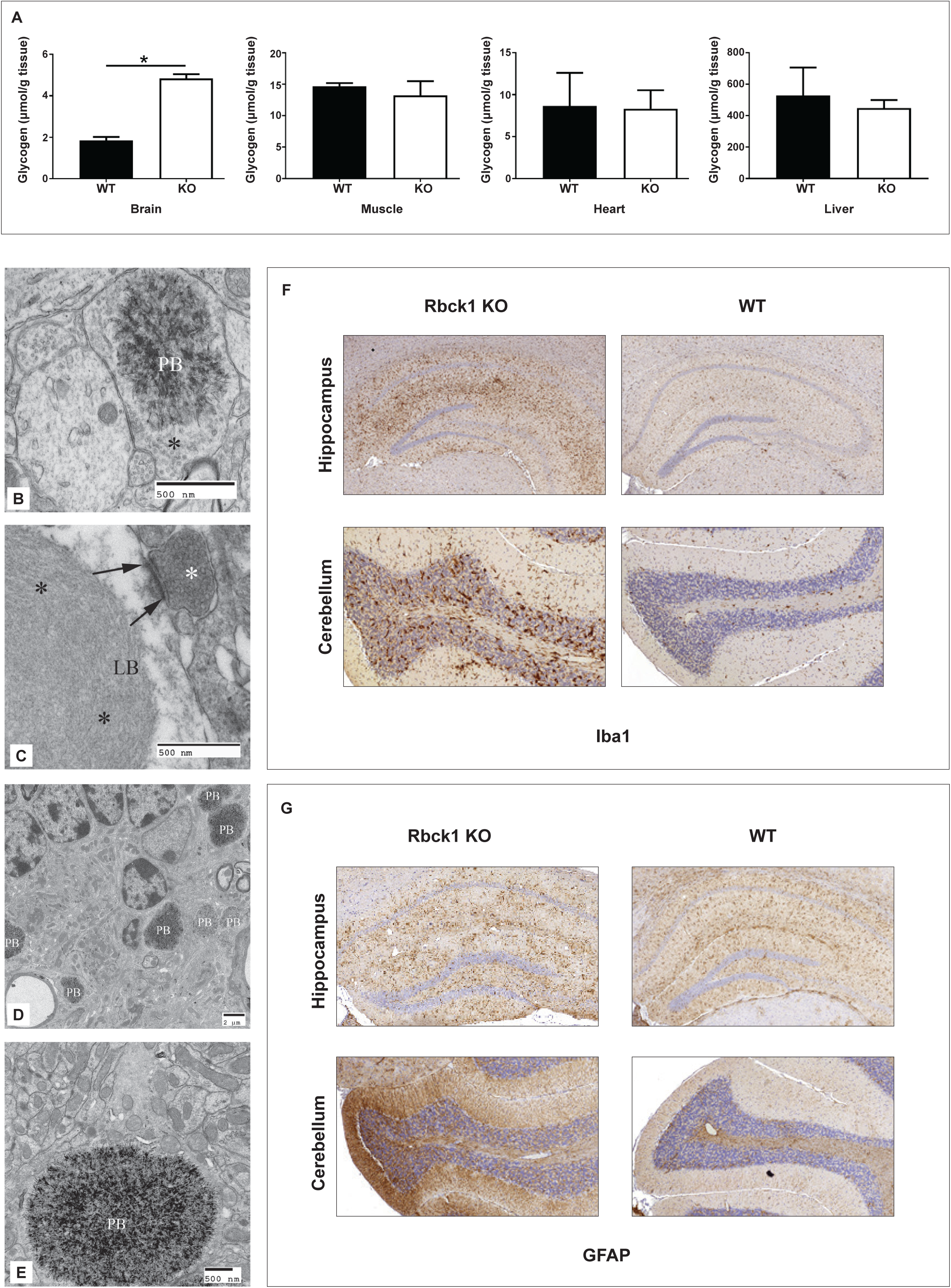
Tissue glycogen content, subneuronal PB location and neurodegeneration in RBCK1 KO mice (all mice 14-month-old). (A) Glycogen content of brain, muscle, heart and liver of WT (black, n =4) and RBCK1 KO (white, n = 4) mice (the accumulated glycogen in RBCK1 KO brain is the PBs). Error bars: SD. (B) Scanning electron micrograph of a representative PB in an RBCK1 mouse in an axon (note synaptic vesicles indicated by the asterisk) at a synapse. (C) Lafora body (LB) in a dendrite in a laforin KO mouse; asterisk, synaptic vesicles; arrows, postsynaptic density. (D) Section of the cerebellum in an RBCK1 KO mouse; note the numerous PBs. (E) PB in a dendrite in an RBCK1 KO mouse (note the postsynaptic density to the left of the PB). (F-G) Sections of WT and RBCK1 KO mouse brains, immunohistochemically stained for markers associated with neurodegeneration; (F), Anti-Iba1 for activated microglia; (G), Anti-GFAP for astrocytes.

Using scanning electron microscopy, RBCK1 KO PBs could only be demonstrated in neurons and not in glia. Within neurons, they were in the cell soma, usually in a juxtanuclear location. PBs could be found anywhere along the neuronal spines. They were completely absent from myelinated long axons but abundantly present at synapses, where they were found both in the dendritic and axonal cytoplasms of synapses (Figure 3B-E).

Neurodegeneration results in astrocyte activation and astrogliosis^39,40^. Glial fibrillary acidic protein (GFAP) staining revealed marked astrogliosis. Ionized calcium binding adaptor molecule 1 (Iba1) staining, microglial activation, was also markedly increased in hippocampi and cerebellums of 14-month-old RBCK1 KO mice (Figure 3F-G). More specifically in the cerebellum, the most dramatic increase in GFAP and Iba1 is detected in the molecular layer and the granular layer of the simple lobule, respectively.

### Motor coordination is impaired in RBCK1 KO mice

The physical performance of mice was analyzed using a number of phenotypic tests, including balance beam, grip strength, rotarod, open field, vertical pole and gait analysis. While RBCK1 KO mice showed equal levels of lateral and vertical movement in the open field test compared to controls, and equal levels of forelimb grip strength, they performed significantly poorer on the rotarod, balance beam and vertical pole tests (Figure 4A), indicating poor motor coordination. The coordination of these mice was also tested using gait analysis. The gait analysis Regularity Index (RI) is used to determine a mouse’s forelimb-hindlimb coordination while walking^41,42^. This parameter measures the proportion of sets, each containing four steps that include all four feet. The higher the RI, the more coordinated the walking pattern of the mouse, with a lower RI being indicative of more missteps. As seen in Figure 4A, RBCK1 KO mice have a significantly lower RI compared to WT.

**Figure 4.**
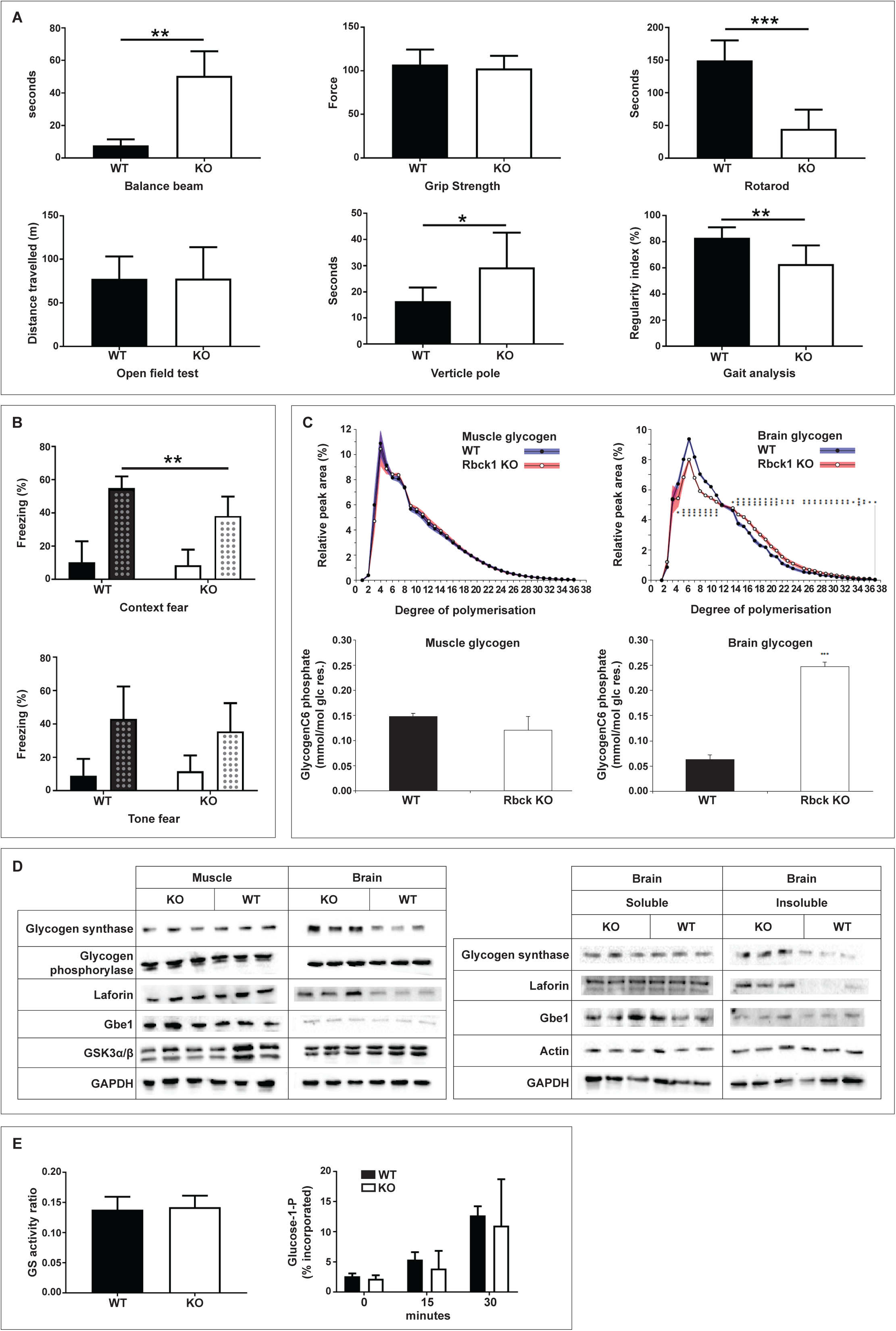
Behavioral testing, glycogen chain length distribution, glycogen glucosyl carbon 6 (C6) phosphorylation level and enzymes of glycogen metabolism in RBCK1 KO mice at 14 months of age. (A) Behavioral testing using indicated procedures; RBCK1 KO (white, n = 6) and WT (black, n = 7) mice. (B) Fear conditioning; RBCK1 KO (white, n = 9) and WT (black, n = 7). Both the context fear (left) and tone fear (right) are presented as the % of time the mouse spent not moving (freezing %). The block pattern represents the baseline level of freezing, the dotted pattern represents the fear response. Error bars: SD. (C) Top panels: chain length distribution of muscle and brain glycogen of RBCK1 KO (black circles, red shade: SD, n = 4) and WT (white circles, blue shade: SD, n = 4) mice. Bottom panels: C6 phosphate content of muscle and brain glycogen of RBCK1 KO (white) and WT (black) mice (error bars: SEM, n = 4). (D) Western blots of enzymes of glycogen metabolism; glycogen synthase, glycogen phosphorylase, laforin, glycogen branching enzyme (GBE1), glycogen synthase kinase (GSK3α/β); RBCK1 KO (n = 3) and WT (n = 3). (E) GS activity ratio and GBE1 activity in brain tissue of WT (black, n = 3) and RBCK1 KO (white, n = 3). Error bars: SD.

### RBCK1 KO impacts learning and memory

Pavlovian fear conditioning was used to analyze aversive learning and memory^43^. Mice are placed in a novel chamber and are subject to hearing a tone, while receiving a foot-shock. The mice are then tested the subsequent day to see how they react to being exposed to the same chamber (context fear) and to the same tone in a different chamber (tone fear). The measurement for fear is the amount of time the mouse stays still (freezing)^44,45^. Importantly, the memory for context and tone represent different brain functions. While damage to the amygdala results in a decrease in the conditioning for both context and tone, lesions of the hippocampus only interfere with the more complex event of the context^46^. As can be seen in Figure 4B, the mean context fear was diminished in the RBCK1 KO mice compared to the controls, with a repeated two-way ANOVA revealing an effect of genotype (F_(1/14)_ = 5.443, p = 0.0351) and time (F_(1/14)_ = 116.3, p < 0.0001) and an interaction of genotype × time (F_(1/14)_ = 4.771, p = 0.0465). Here, time refers to the two conditions, the baseline reading and the post-aversive stimuli reading. Bonferroni’s post hoc test reveals a significant difference between RBCK1 KO and WT in the context fear freezing % (p = 0.007). There was no significant genotype difference for the tone fear (F_(1/14)_ = 0.1647, p = 0.6910), however there was still a time effect (F_(1/14)_ = 46.11, p < 0.0001). There was also no interaction of genotype × time (F_(1/14)_ = 1.423, p = 0.2528).

### RBCK1 KO mice accumulate abnormal glycogen in the brain

The chain length distribution and phosphorylation at glucosyl carbon 6 (C6) of glycogen from RBCK1 KO and WT brain and muscle tissue is given in Figure 4C. The very limited polyglucosan accumulation in the skeletal muscle of these RBCK1 KO mice (Figure 1 and 3A) is consistent with the chain length distribution and C6 phosphate content being indistinguishable from WT. In the brain however, where there is a large accumulation of PBs, the glycogen has a significantly higher proportion of longer chains than in WT mice and an elevated C6 phosphate content, similar to laforin KO mice^47^.

### Glycogen-related enzymes are unaffected by RBCK1 KO

Western blot analyses were performed for a number of the enzymes of glycogen metabolism (Figure 4D). Protein levels were indistinguishable between RBCK1 KO and WT muscle and brain with the exception of glycogen synthase (GS) and laforin in the brain (Figure 4D). These were increased in the total lysate solely due to their accumulation in the insoluble fraction. The relative amounts of GS and laforin were not altered in the soluble fraction of the whole tissue lysate. These results are similar to what has been observed in LD mice^48^, where GS and laforin are trapped in the insoluble PBs by virtue of their strong carbohydrate binding and accumulate, inactive, with the PBs^29,48,49^. The GS activity ratio and GBE1 activity are given in Figure 4E. GS is regulated by covalent modification (mostly phosphorylation) and allosteric effectors such as glucose 6-phosphate. Measurement of GS activity in the presence of low and saturating concentrations of the allosteric activator glucose 6-phosphate is a measure of the activation state of GS as imposed by GS phosphorylation. The results indicate no change in GS activation as the GS activity ratio in RBCK1 KO brain is very similar to that in WT as well as to those previously published^49^. Branching activity in the brain was measured using a phosphorylase stimulation assay with product separation and visualisation by thin layer chromatography^50^. Despite some variances in the RBCK1 KO group on average the branching activity seems similar to WT. In sum, activities of GS and GBE1, which determine glycogen chain length, are unaltered. The soluble quantities of these enzymes and laforin are also unchanged.

## DISCUSSION

The architecture of glycogen molecules is crucial to glycogen solubility and critically includes the regular introduction of branching points. Insufficient branching results in overly long chains that cause glycogen to precipitate and accumulate, which in the brain are associated with neurodegeneration^17^. Until the LD genes were discovered, glycogen architecture was for many decades thought to merely be a function of balanced activities between GS and GBE1. The precise roles of the LD gene products laforin and malin in proper glycogen construction remain unclear. A number of hypotheses are under investigation^14,17,51^, including the following with which we consider the preponderance of available knowledge most closely agrees. The laforin-malin complex appears to represent a safety mechanism that is triggered when glycogen chains are overextended. Through laforin’s carbohydrate binding module the laforin-malin system is recruited to the region in the glycogen molecule with the elongated chain and inhibits GS in that location through malin-mediated ubiquitination. While the role and origin of the phosphate esters in glycogen are still under investigation, it appears that laforin removes the added phosphate when glycogen chains are reshortened, possibly to allow the chain reshortening, because the glycogen digesting enzyme, glycogen phosphorylase is known not able to digest beyond phosphorylated glucosyl residues^17,47,51^.

Recently, RBCK1 was likewise implicated as mice and patients deficient of the protein were shown to accumulate PBs in muscle tissue, in the case of humans resulting in skeletal and cardio-myopathy. Here we report that in RBCK1 KO mice PBs are much more abundant in brain than in skeletal muscle and heart. The distribution of PBs in the brain closely follows the pattern in LD, which differs from APBD. The PBs are in neurons only, while in APBD they are also in glia^52^. They are in the somatodendritic compartments of neurons and not in myelinated axons, in contrast to APBD where they are primarily in axon hillocks and within myelinated axons^7^. One respect in which they do diverge from PBs of LD is that at synapses PBs are in both axonal and dendritic cytoplasm, while in LD they are only on the dendritic side^53^.

As mentioned, RBCK1’s best established function is that of an auxiliary protein within the LUBAC complex, where the actual ubiquitination is carried out by the RBR domain of HOIP. However, RBCK1 also has an RBR domain and may be capable of ubiquitinating substrates independent of LUBAC. Complete absence of HOIP is incompatible with life based on murine studies^54^, but most recently a patient with a homozygous missense *HOIP* mutation was reported who did exhibit PBs on muscle biopsy^38^. While this suggests that RBCK1’s role in glycogen metabolism is via LUBAC, it must be noted that absence of individual LUBAC proteins is well documented to affect the cellular amounts of other members of the complex^55^, and in the above HOIP patient there was a substantial decrease in cellular amounts of RBCK1^38^. As such, it remains unclear whether RBCK1 acts in glycogen metabolism singly or via LUBAC.

The function of RBCK1 (or LUBAC) in glycogen metabolism is likely similar or related to that of the laforin-malin system and dissimilar from that in GBE1 deficiency. Not only are the brain loci of PB formation much more closely aligned between RBCK1 KO and LD mice than between RBCK1 KO and GBE1-deficient mice, but also RBCK1 KO glycogen has the characterstic signature of LD glycogen, namely hyperphosphorylation. Given that RBCK1 and LUBAC share E3 ligase activity with the LD protein malin, the simplest explanation might be that they play a role similar to that of malin, potentially involving regulating GS activity. Like in LD, this regulation would likely occur solely at the few glycogen molecules that develop overextended branches at any one time. This would be consistent with the absence of an effect on overall GS protein and activation state in tissues of the RBCK1 KO mice (Figure 4D-E) or LD mouse models^49^. PBs in APBD patients and the GBE1-deficient mice result from an ∼80% loss of GBE1 activity^6,56^. GBE1 activity in the RBCK1 KO mice is indistinguishable from that in WT, indicating that RBCK1 deficiency does not cause PBs through an effect on GBE1 activity.

Why PBs remain in the somatodendritic compartment of neurons in LD and RBCK1 KO but seem to travel into and accumulate in long myelinated axons in APBD is unknown. Hyperphosphorylation, the distinguishing feature of LD glycogen, has been suggested as a possible factor^57^, but remains unconfirmed. One remarkable difference between RBCK1 KO and LD PBs is at synapses. While LD PBs are found only in the dendritic side of synapses, RBCK1 KO PBs are present also in the axonal side of synapses. That they appear, like those of LD, not to travel down axons, suggests that in the case of RBCK1 KO they may form locally in axonal termini, and that RBCK1 may therefore have a glycogen-related role in this location that the LD proteins do not.

Patients with *RBCK1* mutations have a hyperinflammation/immune-deficiency disease that ranges from no clinical symptoms to fatality^18,20,28^. While this spectrum was initially considered to relate to gene region specificity of mutations^55^, presently, with larger numbers of patients known, this correlation can no longer be made. For example, there are patients with complete loss-of-function mutations with or without clinically manifested immune disease^58^. The RBCK1 KO mouse (the same used in the present study) is immunologically healthy, unless exposed to particular pathogens, and its immunodeficiency/hyperinflammation phenotype is modified depending on which chronic background virome it harbors^58,59^. All patients with *RBCK1* mutations have the glycogen storage disease and succumb from it to cardiac failure, unless transplanted^18,60^. To date, this newly recognized glycogen storage disease has not been associated with a brain phenotype. It is possible that RBCK1 is less relevant in human compared to mouse brain glycogen metabolism. Alternatively, now cardiac-transplanted RBCK1-deficient patients may yet develop neurological disease.

In conclusion, this data is indicative of a new layer of regulation of glycogen metabolism. The similarities with LD and differences from APBD, together with absence of any alteration of branching enzyme, suggests that this system intersects or parallels that of laforin-malin. Future work, now on both the laforin-malin and the RBCK1 tracks, should elucidate the basic mechanisms of proper three-dimensional construction of the glycogen molecule.

## CONFLICT OF INTEREST

The authors declare no conflict of interest.

## AUTHOR CONTRIBUTIONS

M. A. S. designed and performed experiments, analyzed experimental data and prepared the manuscript (including completing 1^st^ draft). F. N. performed experiments, analyzed experimental data, and helped prepare the manuscript. E. E. C. designed experiments and analyzed experiment data. L. F. D. and M. C. performed experiments and analyzed experimental data. A. M. P. performed experiments, S. M. helped prepare manuscript and X. Z. performed experiments. C. A. A. performed experiments and analyzed experimental data, L. I. performed experiments, S. A. performed experiments and P. W. performed experiments and analyzed experimental data. B. A. M. designed experiments, analyzed experimental data and helped prepare manuscript.

## ACKNOWLEDGEMENTS

We would like to acknowledge the contribution of Igor Vukobradovic in the Clinical Phenotyping Core at The Centre for Phenogenomics for help with behavioural phenotype testing. We also thank Thea Darnell for her help with the design and layout of figures. M.A.S was supported by an National Health and Medical Research Council CJ Martin Fellowship GNT1092451. This work was funded by the Ontario Brain Institute and the National Institute of Neurological Disorders and Stroke of the National Institutes of Health under award number P01 NS097197. Berge A. Minassian holds the University of Texas Southwestern Jimmy Elizabeth Westcott Chair in Pediatric Neurology.

## CONTACT FOR REAGENT AND RESOURCE SHARING

Further information and requests for resources and reagents should be directed to and will be fulfilled by the Lead Contact, Berge A. Minassian.

## EXPERIMENTAL MODEL AND SUBJECT DETAILS

### Animals

Female *Rbck1*^-/-^ (B6.Cg-Rbck1^tm1Kiwa/tm1Kiwa^)^61^, *Epm2a^-/-^* (B6.129-Epm2a ^tm1Kzy/tm1Kzy^)^62^ and *Gbe1^Y329S/Y329S^* (*Gbe1^tm2.1Hoa/tm2.1Hoa^*)^36^ mice and WT littermates were housed in ventilated cages at 21 ± 1 °C, under a 12 h light-dark cycle with *ad libitum* access to a standard commercially available diet and water. Mice were sacrificed by cervical dislocation and harvested brain, skeletal muscle, heart and liver tissues were divided with half being fixed in neutral buffered formalin (10%) and half being snap frozen in liquid nitrogen. Spinal cords were removed and fixed in neutral buffered formalin. RBCK1 mice were sacrificed at 1 month, 6 months and 14 months. and laforin and GBE1 mice were sacrificed at 12-14 months.

All animal procedures were approved by The Centre for Phenogenomics Animal Care Committee.

## METHOD DETAILS

### Histochemistry

Samples were stained with the following: periodic acid-Schiff staining with diastase predigestion (PASD) was used to stain polyglucosan bodies; glial fibrillary acidic protein (GFAP, rabbit polyclonal anti-GFAP, BioLegend) staining and ionized calcium binding adaptor molecule 1 (Iba1, rabbit polyclonal anti-Iba1, Wako Chemicals) staining were used to assess gliosis.

### Scanning electron microscopy

As previously described^33^, mice were perfused with 2.5% glutaraldehyde in 0.1 M phosphate buffer (pH 7.4) through the heart’s left ventricle. After mincing into ∼1 mm cubes the samples were left to fix for 4 h. The tissue was washed with phosphate buffer and post fixed in 2% OsO_4_ (in phosphate buffer) for 1 h. The tissue was dehydrated in an ascending series of acetone concentrations and were infiltrated, embedded and polymerized in Embed 812-Araldite overnight at 60 °C. Ultrathin slices were cut and stained with uranyl acetate and lead citrate. Scanning electron microscopy was then used to analyze the sections (JEOL JEM 1011, Peabody, MA).

### Behavioural testing

All mice were acclimatized to the behavioural testing room for 30 min prior to testing.

#### Rotarod

An accelerating rotarod (4 - 40 rpm over 300 s) was used to assess motor coordination. Latency to fall or to complete two passive rotations was measured. Each mouse performed four trials with a 30 min inter-trial interval. The longest three of four scores were averaged.

#### Vertical pole

The vertical pole test was performed as previously described^63,64^. Mice were placed on top of a rough-surfaced vertical pole (45 cm high, 1.1 cm in diameter). The time for each mouse to turn downward (t_1_) and the total time to reach the floor (t_2_) were recorded (t_2_ cut-off limit being set to 120 s). Each mouse performed three trials. The difference between t_2_ and t_1_ was averaged across the trials.

#### Balance beam

The time taken for mice to traverse a round balance beam (90 cm long, 18 mm in diameter) suspended 50 cm above the floor was recorded (cut-off limit set to 60 s). Mice underwent four consecutive training trials the day prior to testing.

#### Grip strength

Maximum forelimb grip strength (peak tension) was measured using the Animal Grip Strength System (San Diego Instruments). Mice underwent five consecutive trials and all scores were averaged.

#### Gait analysis

The ExerGait (XL) treadmill (Columbus Instruments) was used to analyze mouse gait. The treadmill speed was set to 19 cm/s and the camera frame rate was set to 100 frames/s with a maximum of 2000 frames taken. Mice were placed onto the non-moving treadmill for 30 s to acclimatize. Recording commenced once the treadmill was turned on and mice reached a constant running pace. The test duration was 20 s, however, if the mouse’s running pace oscillated, the procedure was restarted. The collected video was analyzed using TreadScan software (Clever Sys Inc.).

#### Fear conditioning

The use of Pavlovian fear conditioning is widely used to analyze aversive learning and memory^43^. This method involved exposure to a neutral conditional stimulus (tone), paired with a fear-inducing aversive unconditional stimulus (foot shock); all performed in a novel chamber. After this pairing, the amount of fear exhibited by a mouse, measured as the percentage of time the mouse does not move (freezing percentage), when exposed again to the chamber (contextual fear) and to the tone (cued fear), is indicative of the mouse’s memory. The link between freezing and fear in rodents has been long established^44,45^.

Using an NIR Video Fear Conditioning System for Mouse (Med Associates Inc.), video freeze software was used to score the freezing and movement of mice. Mice were first placed into the novel chamber for 120 s (giving a baseline freezing percentage). This was followed by 30 s of audible tone and 2 s of foot shock (co-terminating with the tone) and then followed by 150 s with no stimuli. After 24 h mice were returned to the same chamber and freezing was recorded for 300 s. This was used to determine the contextual fear (their memory of the environment and of the shock they received the previous day). After 2 h the mice were then placed back in the chamber for 120 s without any stimulus and then 180 s with a second presentation of the tone, measuring the cued fear, again expressed as the percentage of the time when mice froze

#### Open field test

The open field test is used to test for anxiety and exploratory behaviour. A sound attenuating open field arena (43.2 cm^2^) fitted with three 16 beam IR arrays (X, Y and Z axes) and 275 lux LED lights was used with activity monitor software. The open field arena was divided into a peripheral zone (8 cm from the edge of each wall) and a central zone (around 40% of the total area of the arena) and a bright light illuminated the centre zone. The more time a mouse spends in the periphery, away from the light, the more anxious it is considered. Mice were placed in the middle of the apparatus in the dark (lights off). Testing was initiated when the light turned on and the distance travelled, number of rears and percentage of time spent in the centre zone was recorded for 20 min.

### Glycogen extraction

Glycogen was extracted and quantified using a similar procedure to that previously described^65^. Briefly, frozen tissue from brain, skeletal muscle, heart and liver was ground and boiled in 30% [w/v] KOH for 1 h, followed by ethanol precipitation (67% [v/v], 15 mM LiCl) for a minimum of 1 h at −30 °C. Samples were centrifuged at 16 000 g for 20 min at 4 °C, with the supernatants being discarded. The pellets were resolubilized in water by heating samples to 95 °C for 10 min, using intermittent agitation. Ethanol precipitation was repeated three more times, with the final pellet being redissolved in water. Brain glycogen was further purified in order to accurately quantify the C6 phosphate content and the chain length distribution^47^.

### Glycogen content measurements

Glycogen was degraded to glucose using amyloglucosidase using a method similar to that performed previously^66^. Briefly, 5 µL of amyloglucosidase (Megazyme, 3260 Units/mL) was added to a mixture containing 10 µL of glycogen (an aliquot of extracted glycogen redissolved in water), 20 µL of 200 mM sodium acetate buffer (pH 4.5), 2.5 µL of 10% acetic acid and 62.5 µL of deionized water. Two controls with 10 µL of deionized water (instead of sample) were also analyzed. Samples were incubated for 30 min at 50 °C. Samples were then centrifuged at 20 000 g for 20 min, with the supernatant being transferred to a new tube and the pellets being discarded. Using an established glucose determination assay^67^, 5 µL of degraded glycogen and 5 µL of D-glucose standards up to 1 mg/mL was mixed with 170 µL of reaction buffer containing: 150 µL of 200 mM tricine/KOH (pH 8) and 10 mM MgCl_2_; 18 µL of deionized water; 1 µL of 112.5 mM NAPD; 1 µL of 180 mM of adenosine triphosphate (ATP) and 0.5 U of glucose-6-phosphate dehydrogenase (G6PDH; Roche). After the absorbance at 340 nm was recorded for 20 min to determine a baseline, 4 µL of hexokinase solution (0.75 U in 5 µL of 200 mM tricine/KOH [pH 8] and 10 mM MgCl_2_) was added to each well and the absorbance at 340 nm was recorded for 30 min. The concentration of glucose in each well was calculated by subtracting the baseline absorbance from the absorbance plateau, using the D-glucose standard curve. The average glucose concentration of the two water controls was subtracted from each sample, giving a final glucose concentration. This was then used to calculate the glycogen concentration in the initial sample. Samples unreacted with amyloglucosidase were also tested, showing background glucose not coming from glycogen was insignificant and could thus be discounted.

### Chain length distributions

As previously performed^47^, extracted brain and skeletal muscle glycogen (purified further as outlined above) was debranched using isoamylase. Glycogen (10-20 µg) was incubated at 37 °C overnight with 200 U of isoamylase (Sigma), in 110 µL of 10 mM sodium acetate (pH 5). After incubation the samples were heated at 95 °C for 10 min to inactivate the isoamylase and centrifuged at 20 000 g for 10 min at room temperature. A 90 µl aliquot of the supernatant was applied to a High Performance Anion Exchange Chromatography with Pulsed Amperometric Detection (HPAEC-PAD) system (ThermoFisher, ICS5000). The debranched glycogen chains were separated using a CarboPac PA100 column and guard combination (ThermoFisher) at a constant flow rate of 1 mL/min. The two eluents A (150 mm NaOH) and B (150 mM NaOH, 500 mM sodium acetate) were combined for the following elution profile: 15 min equilibration using 95% [v/v] A, 5% [v/v] B; after injection 5 min 95% A, 5% B; a 9 min linear gradient to 70% A, 30% B; 9 min linear gradient to 55% A, 45% B; 37 min linear gradient to 33% A, 67% B; finally 10 min of 100% B. The resultant chromatograms were analyzed using the Dionex Chromeleon Software (V7.2). The relative peak areas were determined for each chain length, with the four replicates being averaged.

### C6 phosphate analysis

The amount of covalently attached phosphate at the C6 position in the glucose monomers was measured using a method previously described^68^. Briefly, brain and skeletal muscle preparations (see above) were hydrolyzed in 0.7 M HCl for 3h at 95 °C, followed by neutralization with 5 M KOH. The levels of G6P were determined using an enzymatically cycling assay^68^, using authentic G6P as an external standard. The levels of G6P were normalized to the hydrosylate’s glucose content determined as previously described^67^.

### Western blot analysis

Western blot analysis for various proteins was completed essentially as described previously^29^. Briefly, protein concentration was determined and 20 µg of total protein was loaded and separated on a 10% SDS-PAGE. Proteins were transferred to nitrocellulose membranes, blocked with a 5% solution of non-fat dry milk powder. Subsequently, the membrane was probed with commercial primary antibodies. The antibodies used were glycogen synthase rabbit monoclonal antibody (15B1; Cell Signalling), GBE1 mouse polyclonal antibody (B01P) and laforin monoclonal antibody (M02, clone 6C6; Abnova), glycogen phosphorylase mouse monoclonal antibody (9F5) and glyceraldehyde-3-phosphate dehydrogenase (GAPDH; Santa Cruz Biotechnology), GSK3α/β mouse monoclonal antibody (ThermoFisher Scientific), actin mouse monoclonal antibody (BD Transduction Laboratories). The membrane was further blotted with secondary antibodies conjugated with horseradish peroxidase. Proteins were visualized using a chemiluminscent detection kit (BIO-RAD).

### *In vitro* glycogen synthase activity assay

The GS activity assay was performed in 96-well plates containing 50 µl of reaction volume (50 mM Tris [pH 8], 20 mM EDTA, 25 mM KF, 2 mg/ml glycogen, 8 mM glucose-6-phosphate, 1.65 mM UDP[14C]glucose, 1 mM UDP-glucose) and 75 µg of total brain lysate. The reaction mixture was incubated at 30 °C for 45 min. The reaction was stopped by the addition of 100 µl cold 100% ethanol and the total suspension was transferred into a 96-well filter plate. Vacuum was applied to remove the ethanol. Five washing steps with 150 µl of cold 66% ethanol were applied. The plate was dried in a 37 °C incubator for 30 min. 50 µl of scintillation fluid was added to each well and the plate was read for 14C signal using the TopCount microplate scintillation and luminescence counter (PerkinElmer)^69^.

### *In vitro* GBE1 activity assay

The activity of GBE1 was assayed using a previously described procedure^50^. A 50 µL reaction mixture was prepared containing 2.2 U of phosphorylase a, 1 mM AMP, 0.5 M sodium citrate, 50 mM of α-D-[^14^C (U)]-glucose-1-phosphate (specific activity of ∼300 dpm/nmol) and 150 µL of brain lysate (normalized to 1 mg mL^−1^ protein). The phosphorylase a was added last to initiate the reaction and 1 µL aliquots were quenched at 0, 15 and 30 min by loading onto aluminium TLC plates (silica gel 60, Merck). The TLC plates were left at room temperature to dry. The separation of residual glucose-1-phosphate (G-1-P) from those incorporated into the growing polymer was achieved by linear ascending development for ∼3 h in 3:12:4:4 n-butanol:isopropanol:acetic acid (glacial):H_2_O. After the TLC plates had dried they were developed overnight and the radioactivity was determined. The amount of G-1-P incorporated into the glucose polymer, which remained at the bottom of the TLC plate was then calculated as a % of total G-1-P signal.

## QUANTFICATION AND STATISTICAL ANALYSIS

Normality of data were assessed using the Shapiro-Wilk test. Statistical analysis of data that passed this test was performed using a two-tailed Welch’s unequal variance t-test. Statistical analysis for data that did not pass the Shapiro-Wilk test was performed using Mann-Whitney test. Repeated mtwo-way ANOVA was used for the fear conditioning data, with Bonferroni’s post hoc tests. GraphPad Prism v. 7 was used for all of the statistical analysis. Results are presented as the mean ± SD, unless stated otherwise. Statistical analyses with p > 0.05 was considered non-significant and significant differences labelled as: p < 0.05 (*), p < 0.01 (**), p < 0.001 (***).

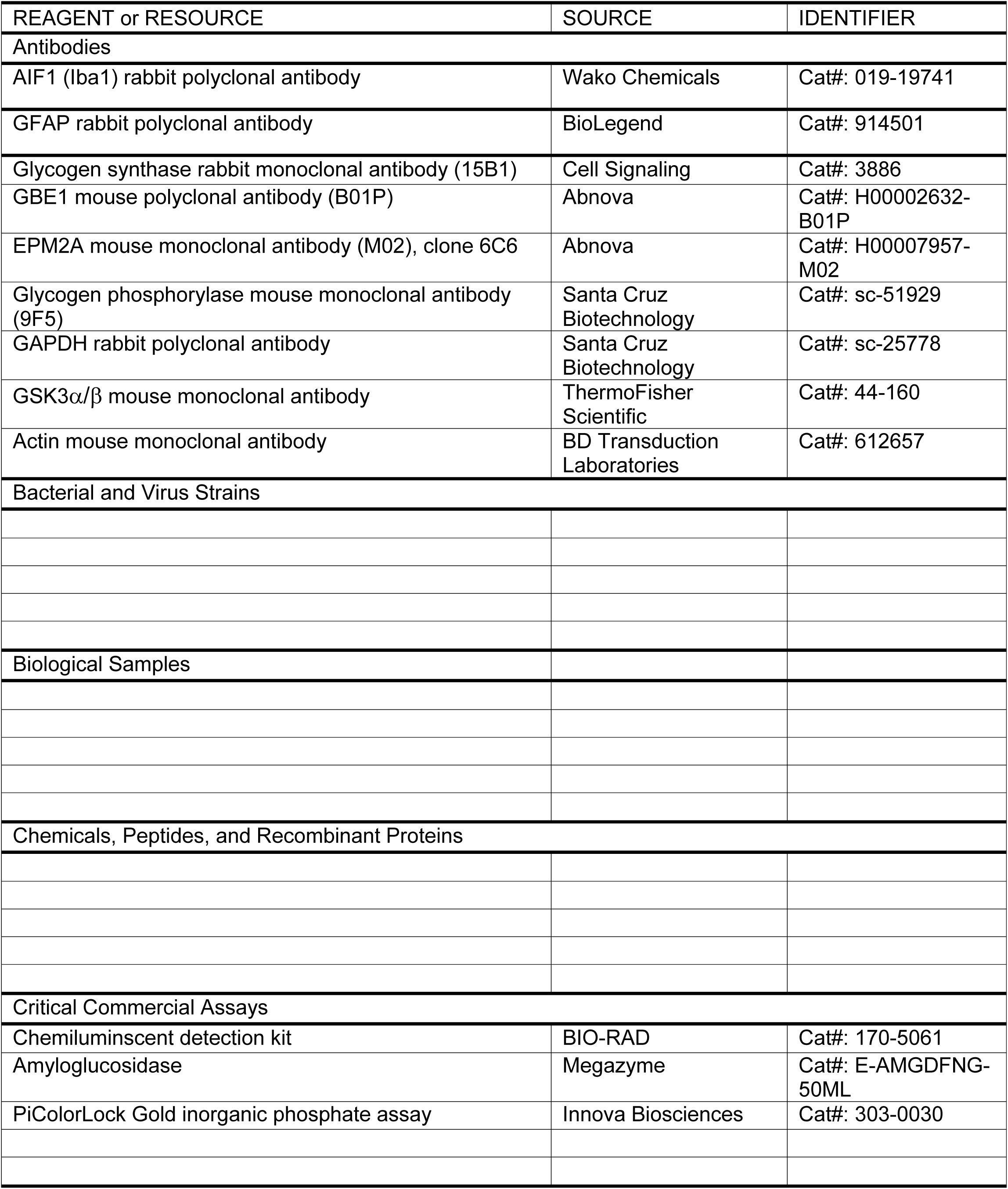

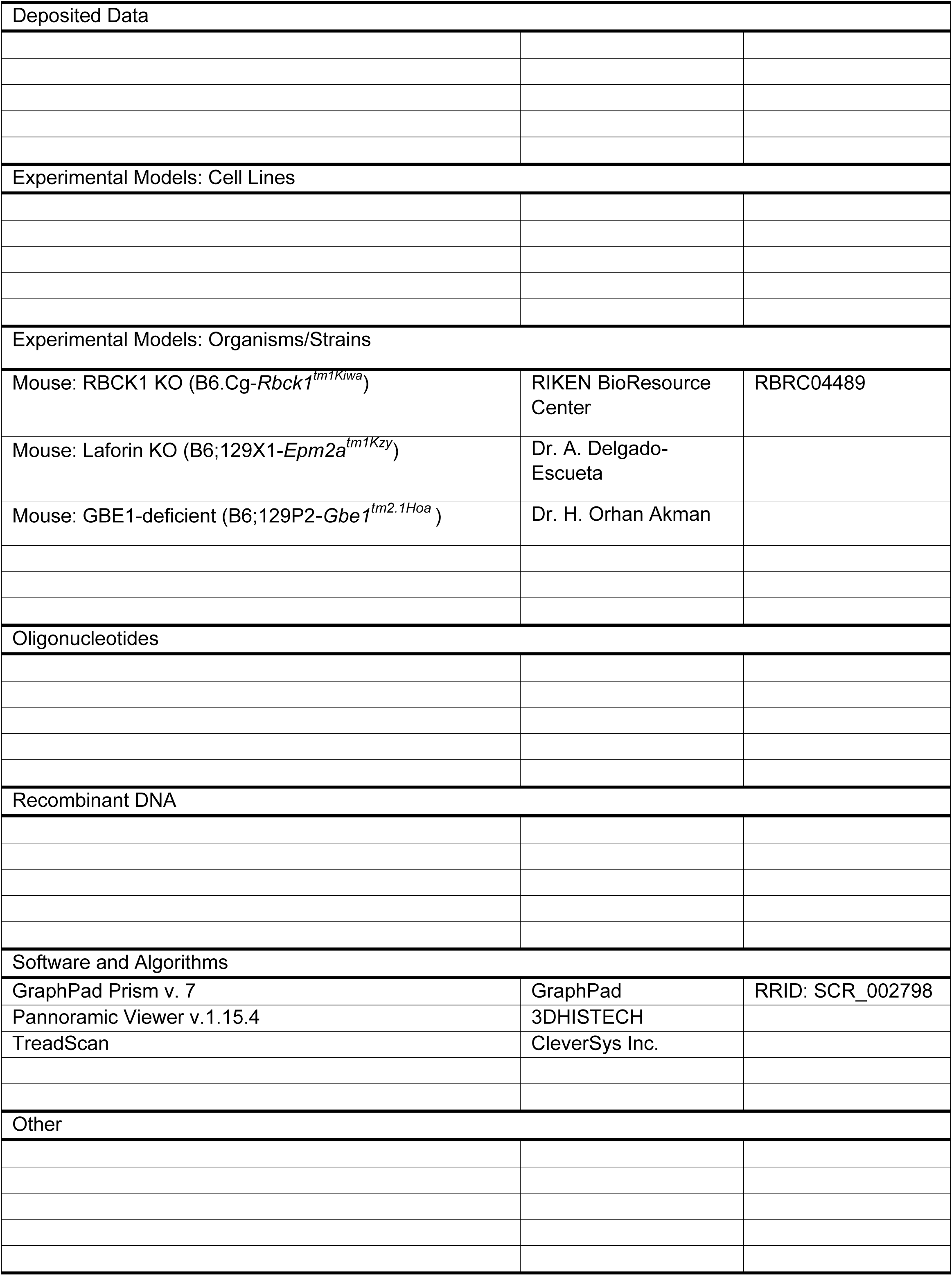

